# Dissociating breathlessness symptoms from mood in asthma

**DOI:** 10.1101/2020.07.15.204289

**Authors:** Olivia K. Harrison, Lucy Marlow, Sarah Finnegan, Ben Ainsworth, Kyle T. S. Pattinson

## Abstract

Asthma is one of many chronic diseases in which discordance between objectively measured pathophysiology and symptom burden is well recognised. Understanding the influences on symptom burden beyond pathophysiology could improve our ability to treat symptoms. While co-morbidities such as anxiety and depression may play a role, the impact of this relationship with symptoms on our ability to perceive bodily sensations (termed ‘interoception’), or even our general and symptom-specific attention is not yet understood. Here we studied 63 individuals with asthma and 30 healthy controls. Alongside physiological tests including spirometry, bronchodilator responsiveness, expired nitric oxide and blood eosinophils, we collected self-reported questionnaires covering affective factors such as anxiety and depression, as well as asthma symptoms and asthma-related quality of life (individuals with asthma only). Participants additionally completed a breathing-related interoception task and two attention tasks designed to measure responsiveness to general temporal/spatial cues and specific asthma-related threatening words. We conducted an exploratory factor analysis across the questionnaires which gave rise to key components of ‘Mood’ and ‘Symptoms’, and compared these to physiological, interoceptive and attention measures. While no relationships were found between symptoms and physiological measures in asthma alone, negative mood was related to both decreased interoceptive metacognitive sensitivity (‘insight’ into interoceptive performance) and metacognitive bias (confidence in interoceptive decisions), as well as increased effects of spatial orienting cues in both asthma and controls. Furthermore, the relationship between the extent of symptoms and negative mood revealed potential sub-groups within asthma, with those who displayed the most severe symptoms without concurrent negative mood also demonstrating altered physiological, interoceptive and attention measures. Our findings are a step towards understanding how both symptoms and mood are related to our ability to interpret bodily symptoms, and to explore how the balance between mood and symptoms may help us understand the heterogeneity in conditions such as asthma.

## Introduction

Asthma is a common and often debilitating chronic condition that affects millions of people worldwide. Asthma is one of the most frequent chronic diseases, in particular amongst children, and has a global prevalence of approximately 1-18%^1^. There is often a significant discrepancy between the disease severity and the extent of symptom burden^2^. Therefore, in this study we wished to conduct a preliminary investigation into possible influential factors that may contribute to symptom discordance heterogeneity within asthma.

Affective dysfunctions such as increased anxiety and depression are some of the most significant co-morbidities in individuals with asthma^3-7^. It is thought that negative mood states (typical in anxiety and depression) may influence symptom perceptions^5^ through a variety of channels, including assigning wider (non-specific) symptoms to asthma diagnoses^8^, reduced perceptions of asthma control^9^ and altering expectations regarding asthma symptoms^10^. Furthermore, negative mood states have also been associated with reduced measures of lung function in asthma compared to neutral states^11^, and treatment of comorbid anxiety and depression with the norepinephrine-dopamine reuptake inhibitor bupropion has been shown to improve both symptoms and spirometry measures^12^.

Importantly, the interaction between physiological (dys)function and symptom perception depends critically on our ability to accurately interpret afferent sensory information from our body, a process termed ‘interoception’^13-15^. Therefore, the relationship between symptoms and physiology may be altered by our ability to accurately interpret these sensations, assign appropriate confidence to our judgements (metacognition), and/or by simply shifting our attention towards or away from them (Figure 1) ^16^. Furthermore, mood disorders themselves have been associated with differences in interoceptive abilities^15,17-20^ and attention biases^21^ in the general population, while somatic symptom disorder has been associated with reduced interoceptive awareness, heightened attention towards symptoms and negative biases when interpreting bodily sensations^17^. Thus, mood may also alter interoception and/or attention towards sensations – either directly or indirectly through inflation of symptoms (Figure 1) in people with breathlessness.

**Figure 1.**
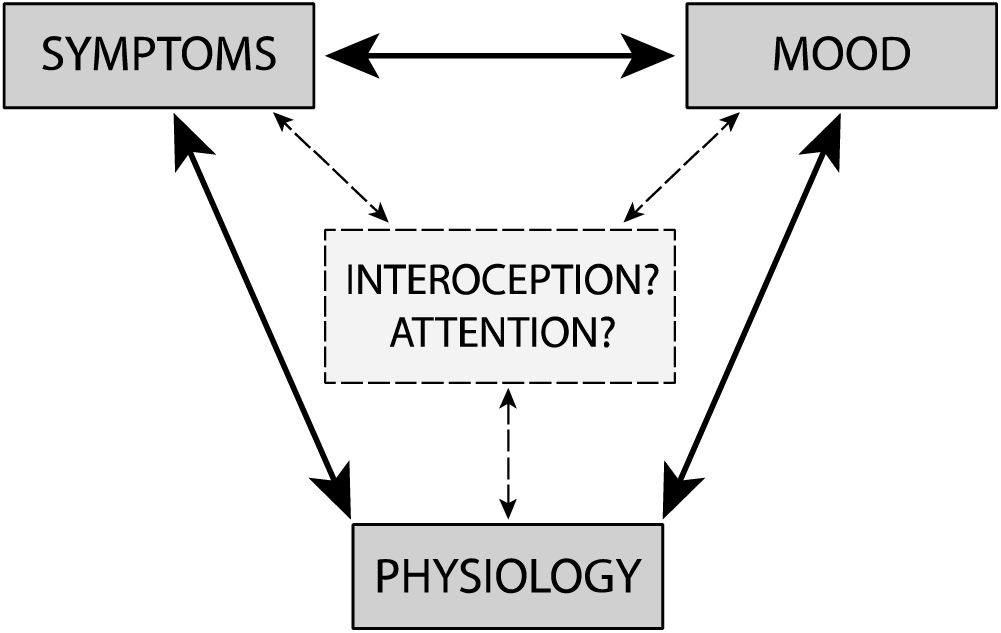
Visualisation of the potential place of interoception and attention within the interaction(s) between symptoms, mood and physiology.

Therefore, alongside physiological measures of asthma, here we assessed the breathing-related interoceptive dimensions. These included a measure of sensitivity towards changes in inspiratory resistance, decision bias (towards over- or under-reporting the presence of a resistance), metacognitive bias (average confidence in decisions regarding the presence/absence of a resistance), and metacognitive sensitivity (correspondence between confidence ratings and performance accuracy) using the Filter Detection Task^22^ in combination with an established model of metacognition^23^. We additionally assessed the effect of both asthma-related fear words using the Visual Dot Probe Task and general spatial and temporal cues on attention using the Attention Network Task) and completed these assessments in both a group of individuals with asthma and healthy controls. General spatial and temporal models of attention identify three functional attentional networks: alerting (activating a vigilant state), orienting (directing cognitive resources towards salient stimuli) and executive control (higher level functions such as resolving conflicting stimuli), measured by the Attention Network Test^24^. Additionally, the Visual Dot Probe Task^25^ examines allocation of attentional resources to affective (i.e. asthma-related) stimuli compared to neutral stimuli.

Therefore the aims of this study were as follows:

1. To dissociate the relationship between symptoms, mood and physiological measures in people with asthma.
2. To determine how mood and symptoms relate to measures of breathing-related interoception, breathing-related attention and general attention in asthma (inner triangle visualised in Figure 1).
3. Assess whether sub-groups of individuals with asthma could be identified based on mood and symptom profiles.

## Methods

### Participants

93 participants (58 female, mean age ± sd: Asthma = 44 ± 12 years, Controls = 44 ± 12 years, range 18 – 65 years) were recruited to the study through recruitment letters sent to patients with asthma from several GP practices and via poster advertisements. 63 participants had a doctor diagnosis of asthma; the remaining 30 participants were healthy with no significant disease or illness (see supplement for full contraindications). Written informed consent was obtained from all participants prior to the start of the study. Study approval was granted by East Midlands - Nottingham 1 Research Ethics Committee (17/EM/0107 ID: 216046).

### Data collection procedures

#### Physiological measures

Weight and height measurements were taken for all participants. Four millilitres of venous blood (whole blood) was acquired from the antecubital fossa by a trained researcher according to University of Oxford and Oxford University Hospitals venesection policy from both healthy volunteers and asthma group volunteers. The level of nitric oxide in exhaled breath was measured using a NIOX Mino device (Healthcare21, Basingstoke UK). Spirometry measurements (FEV1, FVC and peak flow) were collected from participants before and after administration of a bronchodilator. Following the first spirometry assessments, participants received 400 μg of salbutamol and the second session of spirometry measurements were taken 15 minutes following administration as per the European Respiratory Society guidelines^26^. At each spirometry assessment a minimum of three measurements in which the researcher confirmed correct technique were collected and the largest result was chosen for each measure^27^. Participants were instructed to not use short-acting inhaled drugs for 4 hours prior to the testing session and to not use long-acting β-agonist bronchodilators and oral therapy with aminophylline or slow-release β-agonists for 12 hours prior to the testing session^27^. A full medical history, including current medications, was taken from all participants.

#### Questionnaires

The following self-report questionnaires were completed by all participants: State-Trait Anxiety Inventory (STAI)^28^, Anxiety Sensitivity Inventory (ASI)^29^, The Center for Epidemiological Studies Depression Scale Revised (CESD-R-20)^30^, Health Anxiety Inventory (HAI)^31^, Multidimensional Assessment of Interoceptive Awareness (MAIA)^32^, Dyspnoea-12 (D-12) Questionnaire^33^, Nijmegen Questionnaire (NQ)^34^, Fatigue Severity Scale (FSS)^35^. In addition to the above questionnaires, participants with asthma completed the following questionnaires: Catastrophic Thinking Scale in Asthma (CaA)^36^, Beliefs about Medicines Questionnaire (asthma) (BMQ)^37^, Medication Adherence Scale (MAS)^38,39^, Asthma Control Test (ACT)^40^ and Asthma Quality of Life Questionnaire (mini-AQLQ)^41^. All questionnaires were scored according to their respective manuals and descriptions of each questionnaire can be found in the supplementary materials.

#### Interoceptive Filter Detection Task

Participants completed a respiratory resistance detection task based upon the protocol used by Garfinkel et al.^42^. Participants were asked to breathe through a breathing circuit (Figure 2) and following a cue from the researcher determine if a respiratory resistance was added, reporting their response and confidence in their decision. To complete this task, participants breathed through a disposable anti-bacterial and anti-viral mouthpiece connected to a Hans Rudolph t-shaped inspiratory valve. Via two metres of tubing, this was connected to a set of ultra-low resistance baseline filters. Further filters could be added to the system to increase resistance (minimum 1 filter and maximum 7 filters), or alternatively one filter with the mesh removed functioned as the dummy filter. Full details of the equipment used can be obtained from the Filter Detection Task (FDT) toolbox (https://github.com/ofaull/FDT).

**Figure 2.**
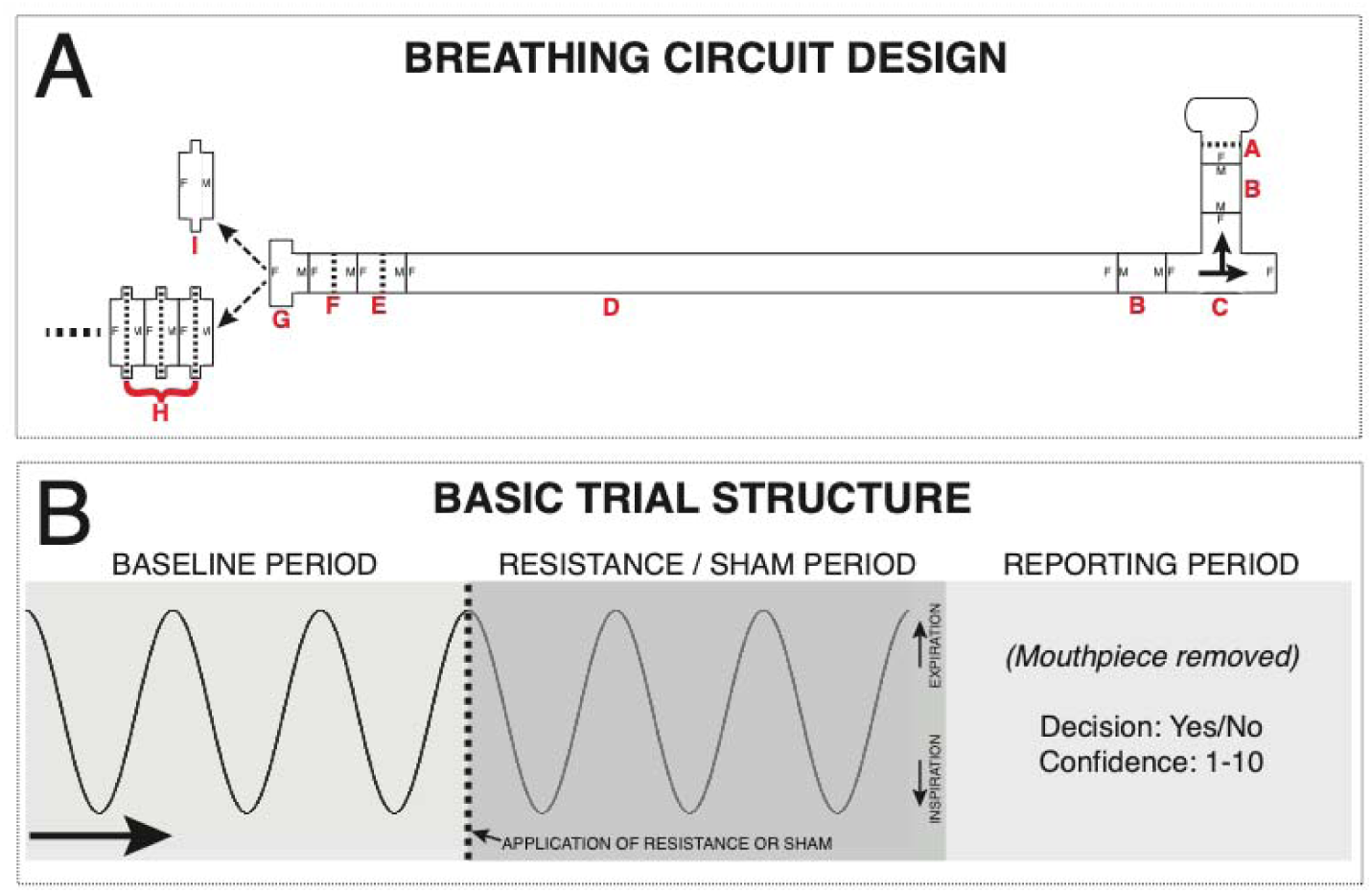
A) Diagram of circuitry for the filter detection task. A single-use, bacterial and viral mouthpiece (A: PowerBreathe International Ltd., Warwickshire, UK - Product SKU PBF03) is attached to a 22 mm diameter connector (B: Intersurgical Ltd., Berkshire, UK - Product 1960000) and a t-shaped inspiratory valve (C: Hans Rudolf, Kansas City, MO, USA - Product 1410/112622), connected to a 2 m length of 22 mm diameter flexible tubing (D: Intersurgical Ltd. - Product 1573000) and two additional baseline filters (E: Intersurgical Ltd. - Product 1541000, and F: GVS, Lancashire, UK - Product 4222/03BAUA). A 22-30 mm (G: Intersugical Ltd. - Product 197100) adapter then allows the attachment of either a series of connected spirometry filters (H: GVS - Product 2800/17BAUF, Resistance at 30 L/min = 0.3 cm H_2_O) or a sham ‘dummy’ filter – a spirometry filter shell with the inner bacterial protection pad removed (I). B) Overview of the basic trial structure for a Yes/No formulation of the task. Participants take three normal size/pace breaths (with the sham filter attached), and during the third exhalation (indicated by the participant raising their hand and the dotted line in panel B) the experimenter either swaps the sham for a number of stacked filters (to provide a very small inspiratory resistance) or removes and replaces the sham filter. Following three more breaths, the participant removes the mouthpiece and reports whether they thought it a resistance was added (‘Yes’) or not (‘No’), and how confident they are in their decision on any scale (here 1-10 used, with 1 = guessing and 10 = maximally confident in their decision). If a two-interval forced choice (2IFC) formulation of the task is used, the filters (resistance) are either placed on the circuit for the first three breaths or the second three breaths according to the FDT algorithm, with the sham filter on the system during the alternate period. The reported decision from the participant is whether they thought the resistance was on in either the first set or the second set of three breaths, and also again the confidence in their decision. Figure taken from Harrison et al. ^22^.

On each trial within the task, participants were asked to take three normal breaths on the system at baseline where a single dummy filter was attached to the system. On their final exhalation at baseline, the participant would raise their hand to indicate that they had completed the baseline breaths. Once the participant raised their hand, the researcher either replaced the dummy with a replacement dummy or a number of real filters. Participants then took 3 further breaths on the system to determine if a resistance had been added or not. At the end of each trial (6 breaths in total), participants were asked to decide whether they thought a resistance had been added to the system (‘yes’) or not (‘no’). Participants were then further asked to specify their confidence in their decision on a scale of 0-100, where a complete guess was rated 0 and complete confidence was rated 100. Participants did not receive feedback on their accuracy/performance.

To complete the task, two or more practice trials were first completed following participant instructions. Task trials were then completed in blocks of 10. In each block, half the trials used a dummy filter on the test breaths and the other half used the test filters. For all participants, the first block of 10 trials compared a dummy filter to 4 resistance filters. To find a threshold for performance, the researcher aimed to find the filter level at which the participant performed at ∼70% accuracy. Accuracy was calculated at the end of each block, if accuracy was below 60% correct, the number of filters used in the next block increased by one filter. If accuracy was above 80% correct, a resistance filter was removed for the next block. To complete the task, the aim was to complete 40-60 trials with a consistent number of resistance filters, where performance was held to between 60-80%.

#### Attention tasks

Participants completed two computer-based tasks to assess attention: The Attention Network Task (Figure 3) and the Visual Dot Probe Task (Figure 4). Stimuli were presented using Inquisit 2 computer software and in both tasks, and participants responded to the direction of a target arrow by pressing the appropriate arrow key on the keyboard. Participants were asked to place their left index finger on the left arrow key and their right index finger on the right arrow key. Instructions for both tasks were presented visually on the screen prior to a practice session. Following each practice session, participants could clarify instructions and the researcher had opportunity to ensure understanding of the task.

**Figure 3.**
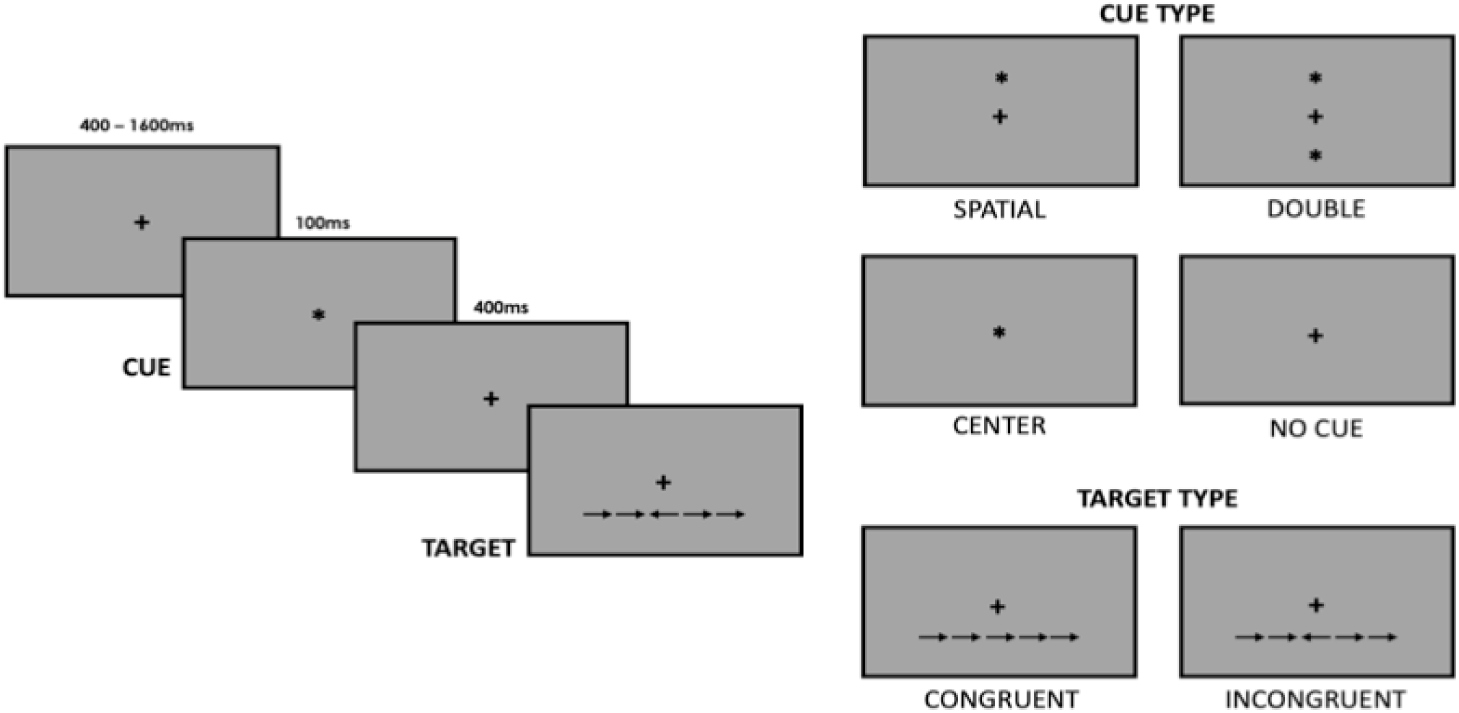
Visualisation of the Attention Network Task (ANT). Participants viewed a central fixation cross for 400-1600ms, followed by a cue (*) or no cue for 100ms. Four cue conditions were used: a ‘centre cue’ where the cue replaced the fixation cross in the centre of the screen; a ‘double cue’ where two cues were presented above and below the fixation position; a ‘spatial cue’ where the cue was presented above or below the fixation position and was indicative of target stimulus location; or no cue was utilised. Following a 400ms intermission (with the fixation cross still present), the target arrow plus two pairs of flankers was presented above or below the fixation cross. The flanker arrows were either congruent or incongruent with the direction of the target arrow. Participants were instructed to respond as quickly and as accurately as possible to the direction of the central arrow using right and left tabs on the keyboard.

**Figure 4.**
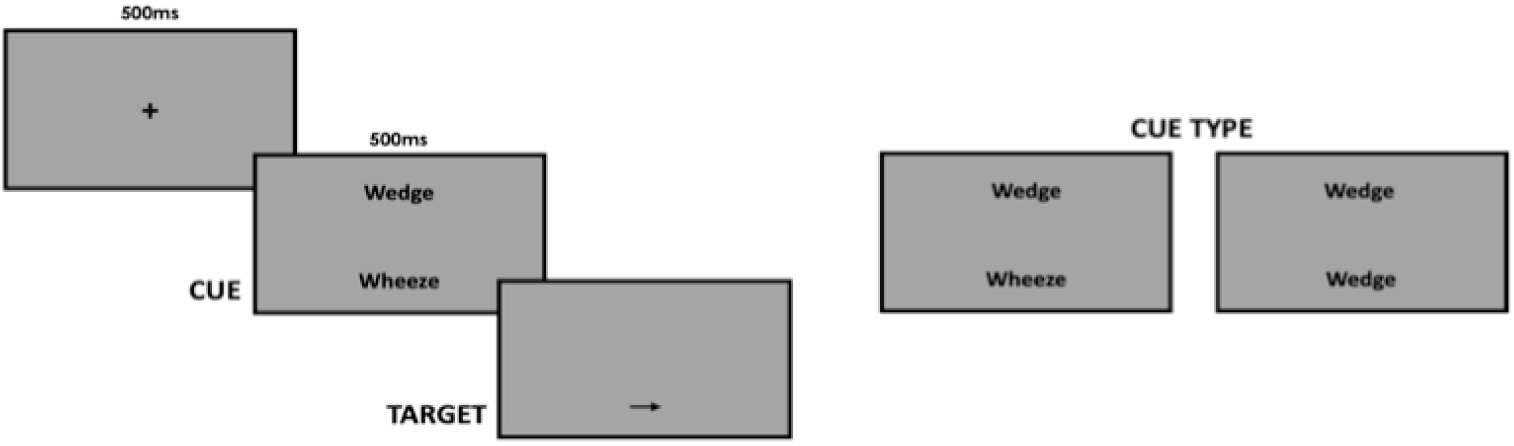
The Visual Dot Probe Task (VPT) utilised a set of asthma fear words and neutral words matched for word length and language frequency to assess attentional bias. A central fixation cross was presented for 500ms before being replaced by two words sitting above and below the fixation location. Two neutral words, or a neutral word and an asthma fear word, were presented for 500ms after which both words were removed with one being replaced by a single arrow stimulus. Participants were required to respond as quickly and as accurately as possible to the direction of the arrow using the left and right arrow keys on the keyboard.

### Calculation of summary measures

#### Physiological measures

Predicted FEV1 and FVC values for each participant were calculated in line with Global Initiative for Chronic Obstructive Disease (GOLD) guidelines^43^. Bronchodilator responsiveness was calculated as the percentage change in FEV1/FVC following administration of salbutamol^44^. A full blood count analysis was conducted for the purpose of measuring blood eosinophils, which are a marker of asthma severity.

#### Questionnaires

Questionnaires were first scored according to their respective manuals. A full correlation matrix was then calculated for (z-scored) questionnaires from the participants with asthma, using MATLAB 2017b (Mathworks, Natick, MA). The structure of the correlation matrices was first examined by applying a hierarchical cluster model to the data (Supplementary Figure 1). Hierarchical models use the covariance across groups of measures in order to organize them spatially within the correlation matrix. The dataset was then visualized in Figure 5 as a connectogram, containing a circular representation of interdependencies between measures. To formalize these relationships between questionnaire measures, an Exploratory Factor Analysis (EFA) was conducted, to uncover latent (hidden) factors. A quality assessment of the EFA model fit was performed by calculating the root mean square residual (RMSR), Tucker-Lewis index (TLI) and the root mean square error of approximation (RMSEA)^45^, and comparing these metrics to established acceptable levels for these indices. Models were fit using Lavaan version 0.6-1 [E22] in R version 3.2.1 (R Core Team). See Supplementary Material for further description of the EFA methods employed.

**Figure 5.**
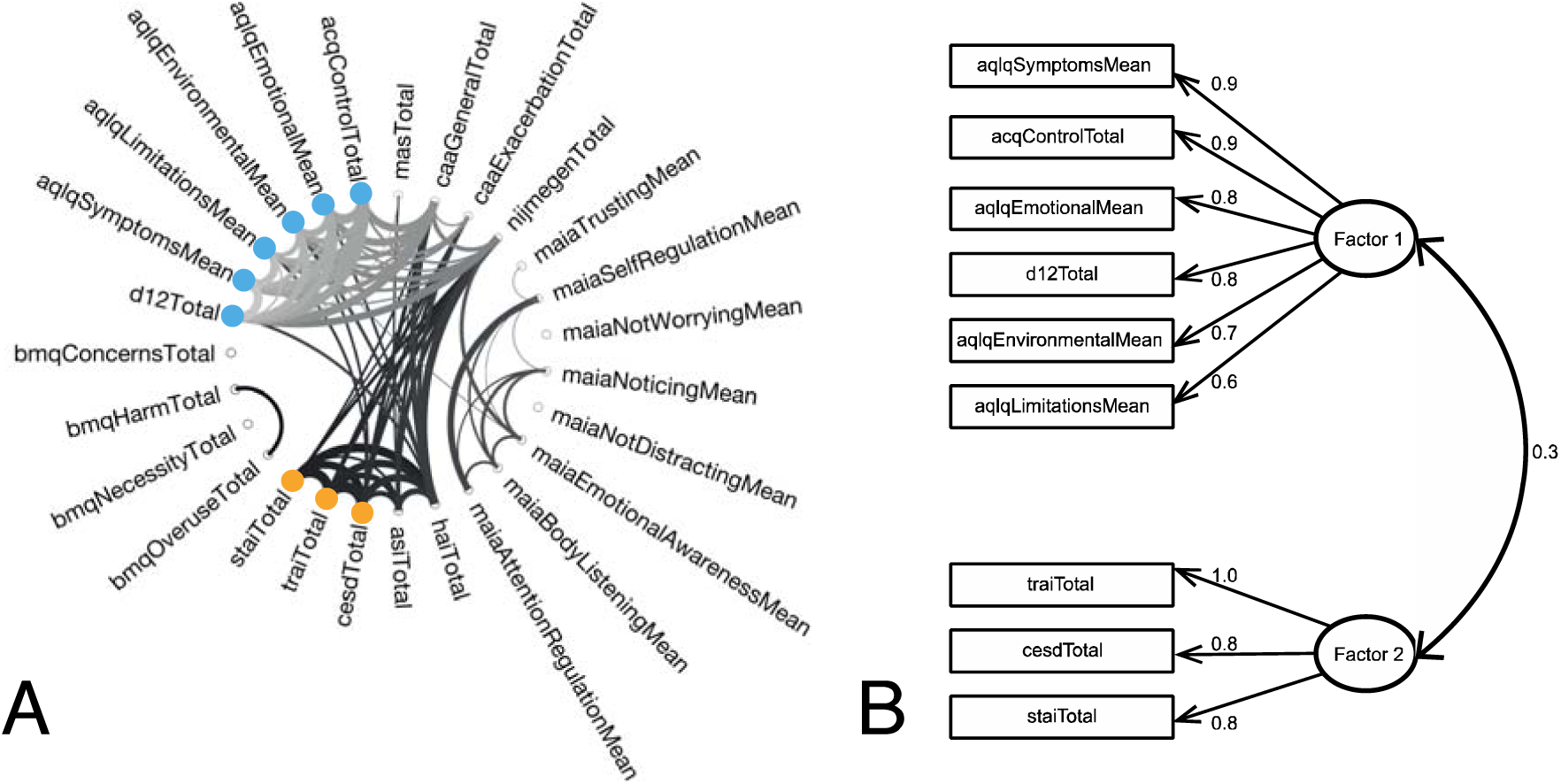
A) Connectogram of all questionnaire metrics collected in the participants with asthma (n = 63). Connections above R>=0.35 are displayed, with line thickness denoting connection strength (colour gradient flows clockwise from light to dark for display purposes only). The two significant latent factors are displayed using blue (Factor 1) and orange (Factor 2). B) Exploratory factor model structure, displaying the two latent factors with their questionnaire loadings and cross-loadings. Questionnaire abbreviations: d12, Dyspnea 12; aqlq, Asthma quality of life; acq, Asthma control questionnaire; mas, Medicines adherence scale; caa, catastrophising asthma; maia, Multidimensional assessment of interoceptive awareness; hai, health anxiety inventory; asi, anxiety sensitivity index; cesd, Centre for epidemiological depression questionnaire; trai, Trait anxiety inventory; stai, State anxiety inventory; bmq, Beliefs about medicines questionnaire.

#### Interoceptive Filter Detection Task

The FDT was analysed using the hierarchical HMeta-d statistical model^23^, with model fits implemented in MATLAB (2017b) and sampling conducted JAGS (v3.4.0). This model firstly utilizes signal detection theory^46^ to provide single subject parameter estimates for task difficulty (d’) and decision bias (*c*), where larger d’ indicates a greater discrimination between stimuli and a negative *c* indicates a bias towards reporting ‘yes’ (over-reporting the presence of a resistance), while a positive *c* denotes a bias towards reporting ‘no’ (under-reporting). Additionally, the model uses a hierarchical Bayesian formulation of metacognitive sensitivity, which is calculated by fitting the ‘metacognitive’ task difficulty parameter meta-d’, normalizing these values by single subject d’ to create estimates of Mratio (meta-d’/d’) that are independent of task performance, then taking the log_e_ of this metric. Metacognitive bias was also calculated as the average confidence score across the analysed trials.

#### Attention tasks

In the Attention Network Task, mean latencies for each participant were calculated for each participant and each attentional condition (central, double, spatial and no cue). The alerting effect was calculated by subtracting the mean double cue reaction time from the mean no cue reaction time. The orienting effect was calculated by subtracting the mean spatial cue from the mean centre cue reaction time. The executive control effect was calculated by subtracting the mean congruent cue reaction time for each cue type from mean incongruent reaction time for each cue type. Incorrect trials and those with a reaction time lying beyond three standard deviations from the participant’s mean were excluded from analysis. Shorter times indicate quicker orienting, alerting and executive control skills. In the Visual Dot Probe Task, breathlessness interference scores were calculated by subtracting the mean response time for threatening asthma words from the reaction time in response to neutral asthma words for each participant (incorrect trials were removed from the analysis). Shorter times indicate quicker reactions induced by threatening asthma words.

### Analysis within asthma

#### Correlations between latent factors and physiology

A correlation matrix was calculated between the factors identified from the questionnaire data and the physiological measures of FEV1/FVC, FEV1 %predicted, bronchodilator responsiveness (% change in FEV1/FVC), exhaled nitric oxide and blood eosinophils. Significance for each of the correlations was set at p < 0.05 with no corrections applied for multiple comparisons.

#### Asthma latent factor regression

A set of linear regressions were then conducted to examine the relationship between the factors identified within the questionnaires and the held-out behavioural scores derived from the FDT (filter number, decision bias, metacognitive bias and metacognitive sensitivity) and the attentional sub-domains (alerting, orienting, executive control and bias). For all except the metacognitive sensitivity analysis, regressions of the EFA scores for the latent factors identified were run against each of the behavioural scores using MATLAB’s fitlm function, with significance was set at p < 0.05 and no corrections applied for multiple comparisons. As the metacognitive sensitivity scores are fit within a hierarchical model, we additionally performed an analogous hierarchical fit of a linear regression using the latent factors scores against interoceptive sensitivity (logMratio) ^22^. Significance for these hierarchical regression coefficients were assessed using one-tailed 95% highest-density intervals (HDIs) on the regression (beta) parameters, to quantify any potential relationships between greater negative behavioural characteristics (such as breathing symptom scores or negative mood) and worsened metacognitive sensitivity (logMratio).

#### Asthma sub-group stratification

To investigate any possible stratification of participants with asthma based on the questionnaire scores, subject-wise clustering was performed on the EFA scores within the asthma group. The most statistically distinct groupings of participants were determined by Matlab’s evalcluster function, which utilises a kmeans clustering algorithm. Each of the groups were then compared to the control group for the physiological, interoceptive and attention measures using either independent t-tests or Wilcoxon rank sum tests (following tests for normal distributions of data), with significance taken at p < 0.05. A single logistic regression model was then applied to the asthma groups using MATLAB’s mnrfit function, with the group that scored the lowest on the symptom and mood factor scores used as the pivot group for comparisons. Significance within the logistic regression was set at p < 0.05, and values are reported as FDR corrected and as exploratory uncorrected results.

Each asthma sub-group was additionally compared to the healthy volunteer group. For all measures except the FDT metacognitive sensitivity, data was tested for normality using the Anderson-Darling test, with an alpha value of p < 0.05 used for rejecting the null hypothesis of normally distributed data. If the data were normally distributed the groups were compared using two-tailed independent t-tests, and if they were not normally distributed non-parametric Wilcoxon rank sum tests were employed. For metacognitive sensitivity scores, frequentist statistics cannot be employed as the values within each group were fit using separate hierarchical models. Therefore, to determine the significance of any group difference in these metacognitive sensitivity (logMratio) estimates, the HDI was calculated across the distribution of sample differences from each of the model fits (as previously described for the HMeta-d model^23^). From this distribution of sample differences, a two-tailed 95% HDI that does not span zero was used to determine any significant difference between the groups^23^. Significance was taken at p < 0.05, uncorrected for multiple comparisons.

### Analysis between asthma and healthy controls

The effect of the latent variables identified were then investigated against the whole cohort of participants (asthma plus healthy controls). However, as the controls did not complete any asthma-specific questionnaires, only the latent mood factor was considered. As the controls were not fit within the EFA model, the first principle component (PC) of the questionnaire scores from the mood factor was calculated across asthma and control groups together. A regression model was utilised that consisted of a group difference regressor, the mood factor and an interaction group*mood regressor, which allows the simultaneous estimation of any difference between the groups, the effect of mood across all participants, and any difference in the effect of mood between the groups (interaction). We then ran this regression analyses on the held-out behavioural measures from the interoceptive and attention tasks, as described above. Significance of regression coefficients was taken at p < 0.05, uncorrected for multiple comparisons. For metacognitive sensitivity, the beta HDIs were calculated across the distribution of samples for each of the three beta estimates in the model, and a two-tailed 95% HDI that does not span zero was used to determine significance^23^. An additional simple group difference analysis between all individuals with asthma and all healthy controls using two-tailed independent t-tests or non-parametric Wilcoxon rank sum tests can be found in the supplementary material.

### Missing data

For all analyses expect the between-group comparisons, missing data were imputed using the Markov chain Monte Carlo method (multiple imputation technique) within the MICE package in R^47^. A summary of the percentage of missing data measures are provided in Supplementary Table 2.

## Results

### Latent factors underlying questionnaire measures in asthma

An exploratory factor analysis performed on the questionnaire measures from the participants with asthma revealed the presence of two underlying latent factors (Figure 5). One of these factors (Factor 1 in Figure 5) consisted of scores of breathlessness symptoms (D12) and asthma quality of life, which we have summarised as an asthma ‘Symptom’ factor, while the other factor included state/trait anxiety and depression scores (Factor 2 in Figure 5), which we have summarised as ‘Mood’. For both factors, increased factor scores represent elevated (worsened) measures of all loading questionnaires. All variables load strongly onto their factors and there is a small amount of correlation across the two factors. The fit statistics of this exploratory structural equation model are shown in Table 1, whereby the RMSR and TLI are within acceptable bounds, while the RMSEA is marginal. The RMSEA is an estimate of the discrepancy between the model and the data per degree of freedom for the model.

**Table 1.**
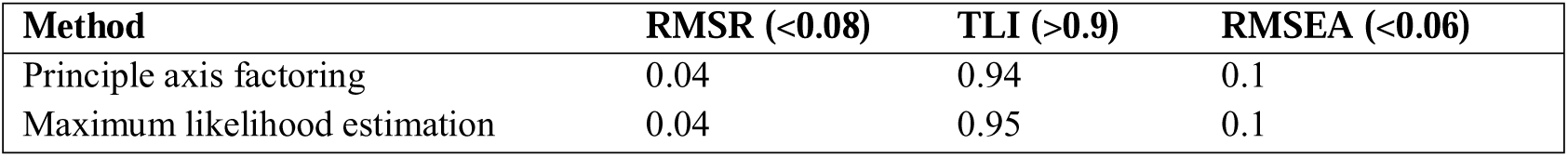
Fit statistics for the exploratory factor analysis model. Abbreviations: RMSR = root mean squared residual, TLI = Tucker-Lewis index, RMSEA = root mean square error of approximation.

| Method | RMSR (<0.08) | TLI (>0.9) | RMSEA (<0.06) |
| --- | --- | --- | --- |
| Principle axis factoring | 0.04 | 0.94 | 0.1 |
| Maximum likelihood estimation | 0.04 | 0.95 | 0.1 |

**Table 2.**
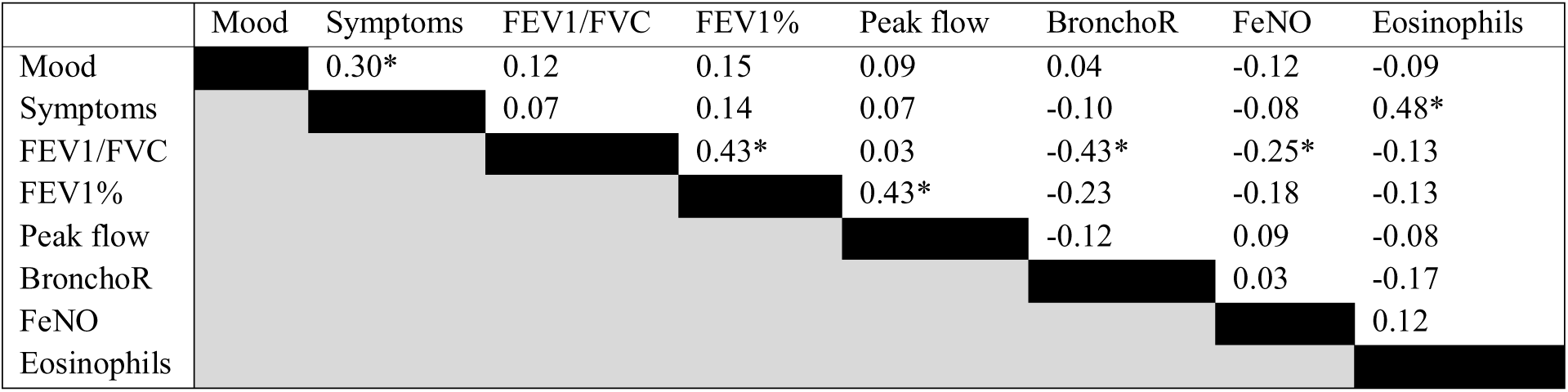
Correlation matrix between latent mood and symptom factors (from questionnaire data) and the physiological measures of FEV1/FVC, percentage predicted FEV1 (FEV1%), peak flow, bronchodilator responsiveness (BronchoR), fraction of exhaled nitric oxide (FeNO) and blood eosinophils. * Denotes significance at p < 0.05.

### Relationship between mood/symptoms and physiology in asthma

A correlation matrix between the two latent questionnaire factors and the four physiological measures within participants with asthma revealed only a significant correlation between symptom scores and blood eosinophils (R = 0.48, p < 0.01), with no physiological measures related to mood scores. Mood and symptom scores were also moderately related (R = 0.30, p < 0.01), as previously reported in the exploratory factor model (Figure 5).

### Relationship between mood/symptoms and interoceptive and attention measures within asthma

A set of exploratory analyses were then conducted where the two latent factors were regressed against all other behavioural measures collected in the FDT and attention tasks. None of these measures demonstrated a relationship with either of the factors, and the results from these analyses can be found in Table 3. No corrections for multiple comparisons were made in these analyses. As noted in the methods section, as the logMratio is fit with a hierarchical statistical model, a ‘significant’ result is denoted when the highest density interval (HDI) does not span zero, signifying that 95% of the posterior distribution lies away from zero.

**Table 3.** Coefficients for the two-factor (mood and symptoms) regression models applied in the individuals with asthma. N.B. Metacognitive sensitivity (logMratio) was fit using a hierarchical regression model combined with a mcmc sampling procedure, and thus the highest density interval (HDI) is presented instead of a p-value. Significance was taken at p < 0.05 (or a 95% HDI that does not span zero) (two-tailed), and no correction for multiple comparison was applied (results are exploratory).

|  | ‘Mood’<br>coefficient | P value<br>(or HDI) | ‘Symptoms’<br>coefficient | P value<br>(or HDI) |
| --- | --- | --- | --- | --- |
| **Filter detection task (FDT)** |  |  |  |  |
| Number of filters (sensitivity) | -0.19 | 0.38 | 0.18 | 0.42 |
| Decision bias | 0.02 | 0.80 | -0.02 | 0.80 |
| Average confidence | -2.43 | 0.28 | -1.30 | 0.56 |
| Metacognitive sensitivity | -0.22 | (-0.54:0.12) | -0.09 | (-0.40:0.20) |
| **Attention network task (ANT)** |  |  |  |  |
| Alerting score | -5.37 | 0.39 | -6.89 | 0.27 |
| Orienting score | -3.87 | 0.56 | -6.95 | 0.30 |
| Executive score | 9.14 | 0.45 | 7.96 | 0.51 |
| **Visual Dot Probe Task (VPT)** |  |  |  |  |
| Breathlessness interference score | 1.08 | 0.83 | -3.14 | 0.54 |

### Sub-group stratification within asthma

Clustering individuals with asthma based on their latent factor scores revealed a three-group structure (Figure 6 and Supplementary Figure 2). Group 1 was characterised by the most severe symptom scores, without any concurrent negative mood scores. Group 2 displayed the mildest symptom and mood scores, while Group 3 demonstrated moderate symptom scores and high negative mood scores (Figure 6). The sub-group scores were compared to healthy controls, where Group 1 was found to have elevated eosinophils yet similar spirometry measures (FEV1/FVC, predicted FEV1, bronchodilation and peak flow) (Figure 7). Beyond the physiological measures, Group 1 also differed in all attention and interoceptive scores when compared to other asthma groups using a logistic regression, however no significant differences were found in these measures when compared to healthy controls (Figure 8). In comparison, Group 2 (mildest symptoms and mood scores, used as the pivot group in the logistic regression) demonstrated the typical decrease in predicted FEV1 compared to healthy controls (Figure 7), and also showed increased metacognitive bias (average confidence) during the interoceptive task (Figure 8). Lastly, Group 3 (moderate symptoms, worst mood) did not demonstrate any significant differences when compared to healthy controls or Group 2 (using logistic regression). A full table of the logistic regression results can be found in the Supplementary Material.

**Figure 6.**
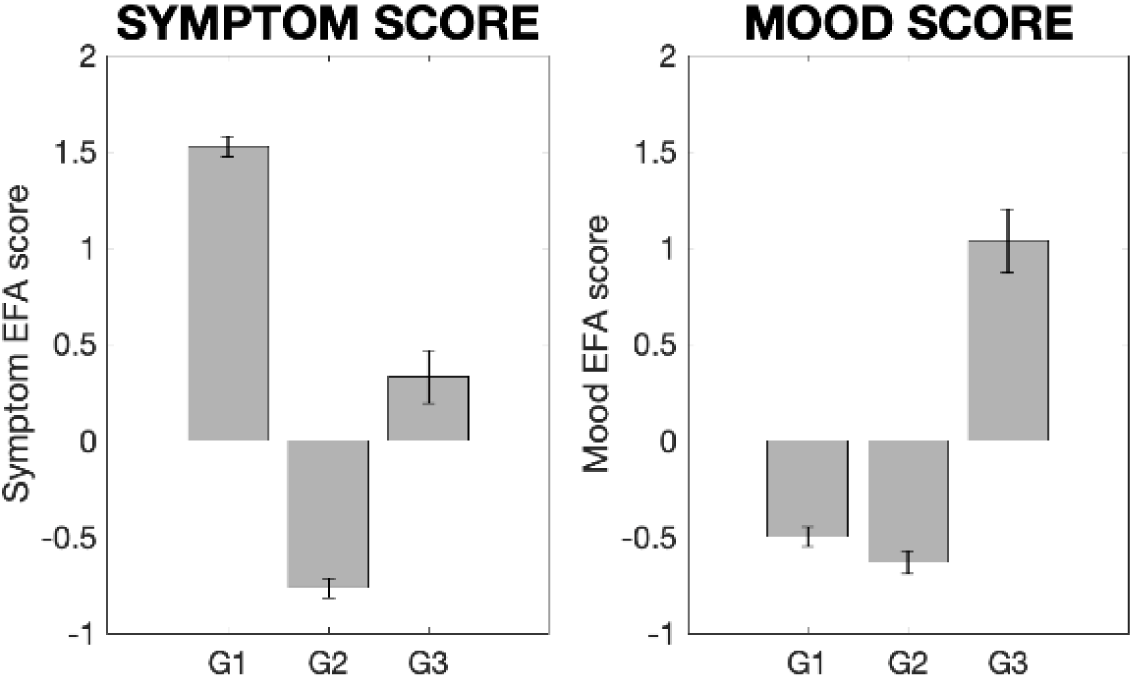
Asthma subgroup scores for the two latent factors identified in the exploratory factor analysis (EFA). Abbreviations: G1-G3, Groups 1-3.

**Figure 7.**
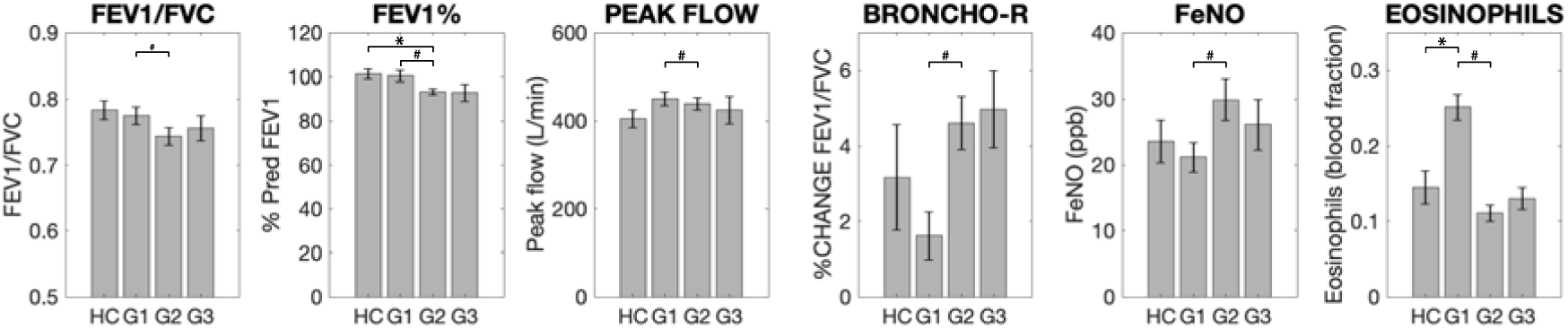
Group means and standard errors of the physiological measures for asthma subgroups (G1-G3) and healthy controls (HC). Each asthma sub-group was compared to healthy controls separately using independent t-tests or Wilcoxon rank sum tests, and the asthma sub-groups were compared using a logistic regression with Group 2 used as the pivot. N.B. Scores FEV1/FVC, percentage predicted FEV1 (FEV1%) and bronchodilator reversibility (broncho-r) were combined using a principle component analysis within the logistic regression due to high correlations. * Significantly different from control group using paired tests (p < 0.05). ^#^ Significantly different from Group 3 using a logistic regression within asthma groups (p < 0.05, FDR corrected for multiple comparisons). Abbreviations: FEV1%, percentage predicted FEV1; BRONCHO-R, bronchodilator responsiveness; FeNO, fraction of exhaled nitric oxide.

**Figure 8.**
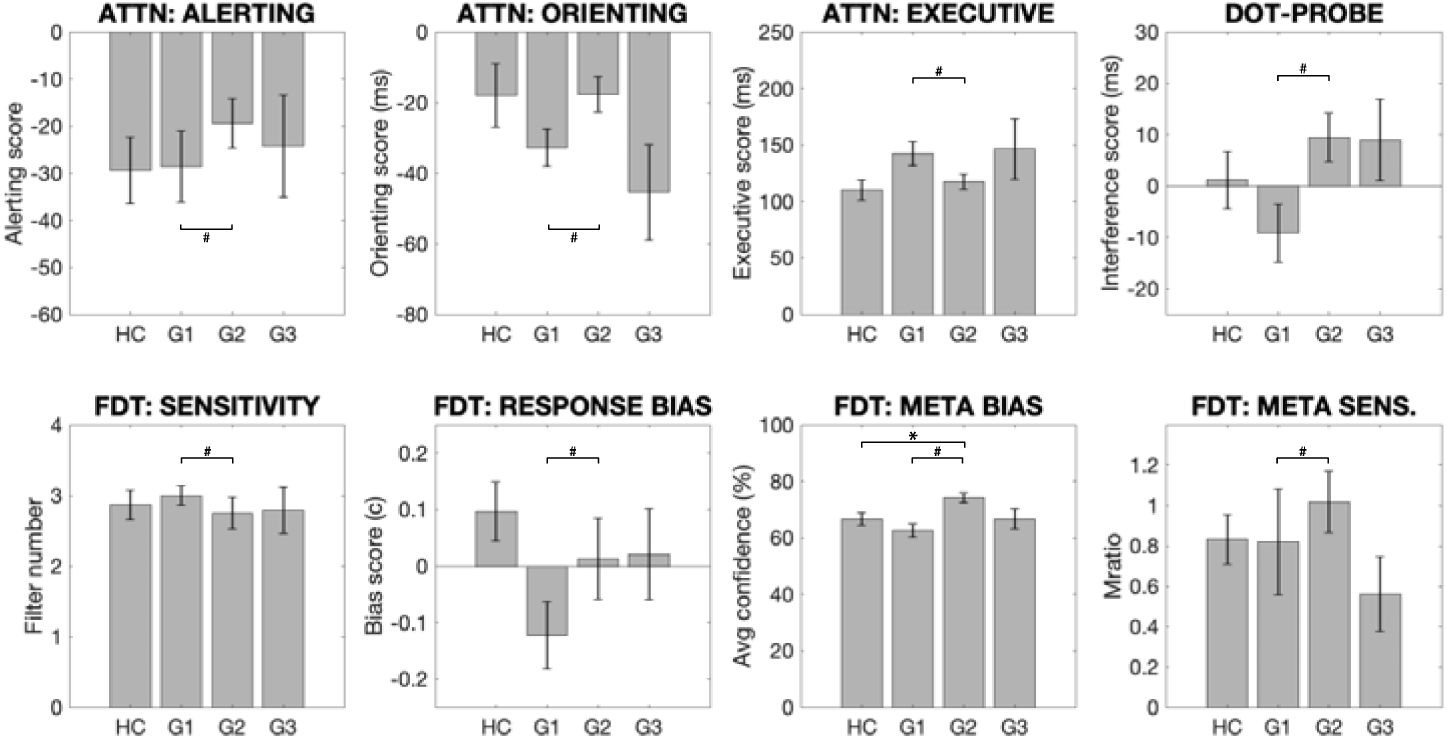
Group means and standard errors of the attention and interoceptive task measures for asthma subgroups (G1-G3) and healthy controls (HC Each asthma sub-group was compared to healthy controls separately using independent t-tests or Wilcoxon rank sum tests, and the asthma sub-groups were compared using a logistic regression with Group 2 used as the pivot. N.B. Scores for metacognitive bias (Meta bias) and metacognitive sensitivity (Meta sens.) were combined using a principle component analysis within the logistic regression due to high correlations. * Significantly different from control group using paired tests (p < 0.05). ^#^ Significantly different from Group 3 using a logistic regression within asthma groups (p < 0.05, FDR corrected for multiple comparisons). Abbreviations: ATTN, attention task; VPT, Visual Dot Probe Task (Breathlessness interference score); FDT, filter detection task; META BIAS, metacognitive bias; META SENS., metacognitive sensitivity.

### Mood factor regression across asthma and healthy controls

A set of regression analyses were then performed across the total cohort of participants, where each of the remaining behavioural measures were regressed against the mood factor scores, an asthma/control group factor, and an interaction between the two (all two-tailed tests). Firstly, a significant effect of the mood factor on logMratio (metacognitive sensitivity) was found across the total cohort of participants, while no effect of group nor any interaction effect was observed (Figure 9A, Table 4). This analysis also revealed a significant effect of the mood factor on average confidence (metacognitive bias) across the total cohort of participants (p < 0.01), and accounting for mood also revealed a marginal group difference (p = 0.05), where individuals with asthma reported higher confidence scores (Figure 9B). However, the interaction effect between mood and group did not reach significance (p = 0.10).

**Table 4.** Coefficients for the regression models (containing an asthma group difference regressor, a mood factor regressor and an interaction term) applied to the total cohort of individuals measured in this study (n = 93). N.B. Metacognitive sensitivity (logMratio) was fit using a hierarchical regression model combined with a mcmc sampling procedure, and thus the highest density interval (HDI) is presented instead of a p-value. * Significance was taken at p < 0.05 (or a 95% HDI that does not span zero (two-tailed)), and no correction for multiple comparison was applied (results are exploratory).

|  | Asthma<br>coefficient | P value<br>(or HDI) | 'Mood'<br>coefficient | P value<br>(or HDI) | Interaction<br>coefficient | P value<br>(or HDI) |
| --- | --- | --- | --- | --- | --- | --- |
| **Filter detection task (FDT)** |  |  |  |  |  |  |
| Number of filters (sensitivity) | -0.05 | 0.79 | <0.01 | 0.99 | -0.13 | 0.58 |
| Decision bias | -0.02 | 0.66 | -0.05 | 0.37 | 0.10 | 0.12 |
| Average confidence | **3.59** | **0.05\*** | **-6.20** | **<0.01\*** | 4.09 | 0.07 |
| Metacognitive sensitivity | 0.13 | (-0.10:0.37) | **-0.36** | **(-0.66:-0.08)\*** | 0.06 | (-0.24:0.38) |
| **Attention network task (ANT)** |  |  |  |  |  |  |
| Alerting score | 1.01 | 0.84 | -0.02 | 1.00 | **-12.50** | **0.05\*** |
| Orienting score | -0.19 | 0.97 | **-15.03** | **0.01\*** | 12.07 | 0.09 |
| Executive score | 10.74 | 0.24 | 3.17 | 0.74 | 7.28 | 0.53 |
| **Visual Dot Probe Task (VPT)** |  |  |  |  |  |  |
| Breathlessness interference score | -0.71 | 0.86 | 7.09 | 0.10 | -8.46 | 0.10 |

**Figure 9.**
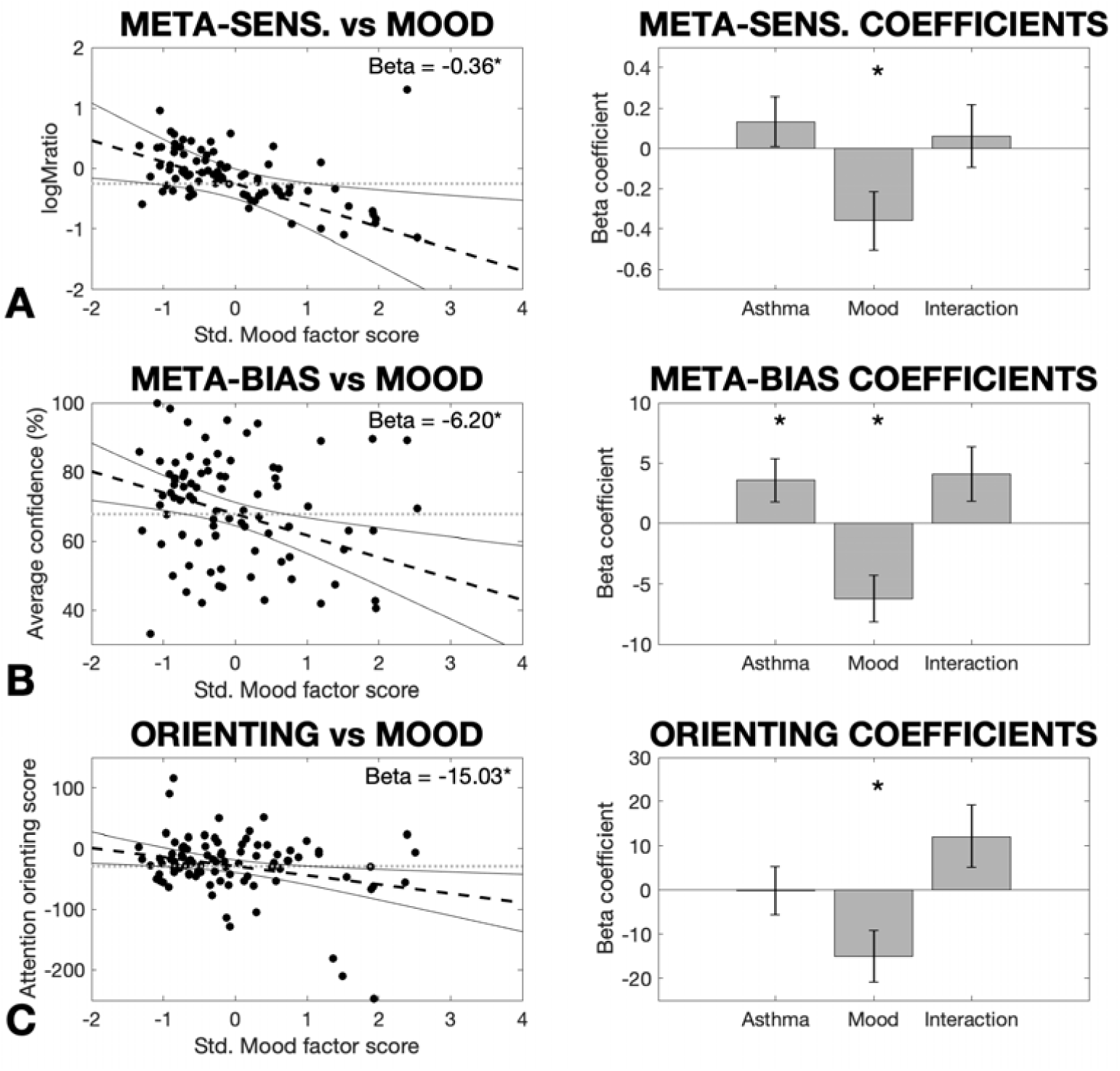
Significant results for the ‘Mood’ latent factor model regressed against the remaining measures: A) logMratio (representing metacognitive sensitivity), B) Average confidence (representing metacognitive bias), and C) Attention orienting. Additional regressors were included for any group difference between asthma and healthy controls, and also an interaction term between group and mood scores. In the left panels, dashed lines signify the regression line, dotted lines are plotted as a comparative ‘null model’ with an intercept term only (zero slope), and solid lines denote the 95% confidence interval of the regression line. The regression confidence intervals visually demonstrate the certainty regarding the estimate of the beta (slope) model parameter, where confidence intervals that do not encompass the ‘null model’ line are required for a significant effect. Note: The logMratio regression was fit within a hierarchical model, which necessitates the use of the confidence intervals for a visual comparison of the uncertainty regarding model beta fits between the hierarchical and standard regression procedures. The average confidence and attention orienting regressions were fit using a standard linear regression model. In the right panels, the mean beta parameter estimate and standard error are shown. * Significantly different from zero (p < 0.05, two-tailed, uncorrected).

For the remaining measures, a significant effect of the mood factor was also found on the attention orienting score across the total cohort of participants (p = 0.01), while there was no significant effect of group (p = 0.97) and the interaction effect did not reach significance (p = 0.09) (Figure 9C). No other measures were found to be related to mood, group or an interaction of the two, with all regression results reported in Table 4.

## Discussion

### Main findings

In this study we firstly separated and characterised the degree of breathing symptoms and negative mood using self-report questionnaire measures, and assessed their relationship to measures of physiology, interoception and attention within asthma. Symptom scores were found to correlate with one physiological measure (blood eosinophils), while negative mood did not relate to any physiological measures. However, using these latent factor scores we revealed preliminary evidence for possible stratification of individuals with asthma into sub-groups, where these groups also demonstrated differences in both physiological, interoceptive and attention scores. Finally, negative mood was related to reduced interoceptive metacognitive sensitivity (or decreased ‘insight’ into breathing-related interoceptive abilities), decreased metacognitive bias (average confidence in interoceptive abilities) and attention orienting across all individuals (asthma and healthy controls), with only metacognitive bias elevated in individuals with asthma compared to healthy controls. These results may guide future studies and hypotheses regarding both the heterogeneity across individuals with asthma and research aimed at developing personalised treatments for breathlessness.

### The relationship between symptoms, mood and asthma physiology

In asthma, the correspondence between the extent of physiological severity and self-report measures of symptom extent is known to be poor^2^. Furthermore, there is a known association between asthma and elevated levels of anxiety and depression^3-7^. However, here we revealed a clear dissociation of specific mood components (i.e. anxiety and depression) from symptom extent (reflected in asthma quality of life scores and dyspnea scores), with only a moderate correlation between these factors. Consistent with much of the literature^2^, symptom scores were only moderately related to one physiological measure of asthma severity (blood eosinophils), while mood scores were not related to any of the physiological measures. While physiology and subjective scores are only weakly related, dissociating breathing symptoms from negative mood allows us to then investigate their independent relationships with important factors such as our ability to perceive bodily sensations (interoception), or our attention towards these sensations.

The variable relationship between symptom and negative mood factors also enabled the identification three sub-groups of individuals with asthma. Within these groupings, those who demonstrated the highest symptoms without a concurrent negative mood state were those individuals with the highest blood percentage of eosinophils, reduced bronchodilator responsiveness and more normalised resting spirometry and expired nitric oxide measures. Therefore, this group may represent those who are least responsive to typical inhaled bronchodilator medications, and thus have a greater level of physiological dysfunction during asthma exacerbations. While the results from these groupings are exploratory due to sample size, the differences between these sub-groups clearly demonstrates that the relationship between symptoms, mood and physiology is complex, and the variability between these measures is likely to contribute to the heterogeneity observed across the spectrum of asthma diagnoses. Furthermore, these results could underlie null effects seen in treatment trials with treatment programs suitable for one sub-group being masked by null effects in other individuals.

### Interoception within the breathing domain

Interoception is an important gateway by which bodily sensations are connected to symptom perception, and here we investigated the relationship between both asthma symptoms and mood and interoceptive domains. Within the breathing-related interoception task (FDT), we firstly found that metacognitive bias (confidence) was related to both an asthma diagnosis and mood scores, while no direct relationships were observed between asthma symptoms and any interoceptive domains. Consistent with previous literature in exteroception^48^, a reduction in confidence regarding interoceptive decisions (metacognitive bias) was significantly related to negative mood across the cohort of both asthma and healthy controls. However, once this mood effect was accounted for, an *elevated* metacognitive bias (i.e. higher confidence scores) was observed within asthma, despite a more negative mood than healthy controls. While the direct interaction effect between metacognitive bias and the asthma group did not reach significance, these exploratory results indicate that there may be a difference in the

confidence assigned to breathing perceptions within asthma. Interestingly, the difference in confidence was most pronounced in those individuals with asthma that reported the most positive mood and the least symptom scores (asthma Group 2). Therefore, it is possible that when exposure to elevated breathing symptoms in asthma is not coupled with worsened mood or subjective symptom burden, an elevation in perceptual confidence (metacognitive bias) can be induced even when absolute interoceptive sensitivity (i.e. the degree of inspiratory resistance that is able to be detected) does not change.

We additionally observed that negative mood was related to reduced metacognitive sensitivity in both asthma and healthy controls – a novel finding within the current metacognitive literature. Metacognitive sensitivity can be considered to reflect insight into one’s interoceptive performance, where an individual is able to more accurately assign greater confidence values on trials when they make correct judgements regarding interoceptive decisions (here the presence/absence of an inspiratory resistance), and lower confidence values when they make incorrect decisions. In previous work oriented towards the external domain (using a visual discrimination task), metacognitive sensitivity appeared to be unaffected by negative mood^48^ despite a decrease in metacognitive bias (overall confidence scores). In contrast, we have demonstrated both a reduction in metacognitive bias *and* sensitivity with negative mood scores within the interoceptive domain, indicating that the effect of negative mood may differentially alter external and internal metacognitive sensory processing. Furthermore, despite an overall more negative mood in asthma, the relationship between mood and metacognitive sensitivity is consistent between asthma and healthy controls, and thus may be a result of general mood factors such as anxiety and depression and independent of the presence of asthma.

Finally, while direct relationships were not identified between interoception and symptoms across the entire asthma cohort, the asthma sub-groups demonstrated important interoceptive differences. In particular, the asthma group with the highest symptoms and elevated blood eosinophils demonstrated a decrease in sensitivity towards detecting inspiratory resistances, a bias towards over-reporting the presence of a resistance, and a decrease in confidence (metacognitive bias) regarding these interoceptive decisions when compared to the low symptom asthma group. Therefore, while the relationship between interoception and symptoms may not be consistent across individuals, here we present evidence that breathing-related interoceptive properties may be disrupted in the presence of elevated asthma symptoms, although the causality of this relationship cannot be determined without a longitudinal intervention that targets these interoceptive abilities.

### General and breathing-related attention

An additionally important aspect in our ability to perceive symptoms from our body is our capacity to attend to stimuli – both in general and in response to symptom-relevant stimuli. While we observed no relationship between either symptoms or mood and attention within asthma, across the total cohort of participants negative mood was associated with improved reaction times as a result of a spatial cue (measured using the attention ‘orienting’ score). Again, no group difference in the attention orienting score was apparent between asthma and healthy controls, indicating that this effect may also be associated with general changes in mood that are independent of asthma diagnosis. However, in a similar vein to the interoception results, the asthma group that displayed the greatest symptoms without a concurrent negative mood state exhibited differences in attention measures. Not only did these individuals have a greater effect of temporal and spatial cues on attention compared to the low symptom asthma group, they also demonstrated a greater bias towards asthma-related fear words in the Visual Dot Probe Task. While these results are exploratory in nature, they provide a platform for future work investigating the potential effect of worsened mood on increased attention towards spatial cues across the population, and also the possibility of altered attention in those who have elevated symptoms in asthma.

### Conclusions

A well-known discordance exists between symptom load and objective measures of physiological dysfunction in asthma, with an elevated prevalence of co-morbidities such as anxiety and depression. Here we conducted preliminary tests to investigate whether both interoception and attention may be important mechanisms by which either symptoms or mood may alter the ability to accurately interpret sensory signals from the body. It appears that mood may directly influence aspects of both metacognition of interoception and general attention – important elements within perceptual pathways – in both asthma and healthy people. Lastly, we were able to utilise the variable relationship between symptoms and negative mood to identify sub-groups of individuals with asthma, who demonstrated distinct differences in physiological, interoceptive and attention measures. While small group sizes limit generalisability of these sub-groupings, we hope that these exploratory results may help generate hypotheses for future studies geared towards understanding the heterogeneity of symptom burden within asthma.

## Acknowledgements

Olivia Harrison (née Faull) is a Marie Sklodowska-Curie Postdoctoral Fellow, supported by the European Union’s Horizon 2020 research and innovation programme under the Grant Agreement No 793580. Kyle Pattinson was supported by the JABBS Foundation, and the NIHR Biomedical Research Centre based at Oxford University Hospitals NHS Trust and The University of Oxford.

## Supplementary Material

### Supplementary Methods

#### Questionnaires

**State-Trait Anxiety Inventory (STAI):** This is 2-part questionnaire with 20 items assessing trait anxiety and 20 items assessing state anxiety. Trait anxiety items are concerned with how the responder “generally” feels. State anxiety items are concerned with how the responder feels “right now” or “at this moment”^28^.

**Anxiety Sensitivity Index (ASI):** This is a 16-item questionnaire which assesses the dispositional tendency to fear the symptoms of anxiety, viewing them as potentially harmful (anxiety sensitivity)^29^.

**The Center for Epidemiological Studies Depression Scale Revised (CESD-R-20):** This is a 20-item questionnaire which investigates depression^30^.

**Health Anxiety Inventory (HAI):** This is an 18-item questionnaire (short version) that assesses anxiety about health independently from health status^31^.

**Multidimensional Assessment of Interoceptive Awareness (MAIA):** This is a 32-item questionnaire that assesses self-report interoceptive awareness across 8 domains: Noticing, Not Distracting, Not Worrying, Attention Regulation, Emotional Awareness, Self-regulation, Body Listening, Trusting^32^.

**Dyspnoea-12 (D-12) Questionnaire:** This is a 12-item questionnaire which is designed to assess breathlessness severity. It has been validated in individuals with respiratory disease^33^. **Nijmegen Questionnaire (NQ):** This questionnaire assesses dysfunctional breathing patterns associated with hyperventilation across 18 items^34^.

**Fatigue Severity Scale (FSS):** This is a 9-item questionnaire that assesses participant fatigue which has been associated with respiratory disease^35^.

**Catastrophic Thinking Scale in Asthma (CaA):** This is a 13-item questionnaire adapted from the Pain Catastrophising Scale that assesses catastrophic thinking during an exacerbation (exacerbation scale) and in general (general scale)^36^.

**Beliefs about Medicines Questionnaire (Asthma) (BMQ):** This is an 18-item questionnaire that assesses specific concerns and representations of medications prescribed to the individual (asthma specific) and general representations of medications^37^.

**Medication Adherence Scale (MAS):** This is a 5-item questionnaire which assesses patient adherence to prescribed medication use for their asthma^38,39^.

**Asthma Control Test (ACT):** This is a 5-item questionnaire that assesses how well the individual’s asthma is managed and controlled over a period of 4-weeks^40^.

**Asthma Quality of Life Questionnaire (mini-AQLQ):** This is a 15-item questionnaire that assesses the impact of asthma on an individual’s quality of life across 4 domains (Symptoms, Activities, Emotions and Environment)^41^.

#### Cluster Analysis

Hierarchical cluster models reorder variables based on their correlation strengths so that groups of related measures sit closer to each other than non-related measures. This allows natural relationships to be easily visualised. The modelling process formalises not only the relationship between pairs of variables, but also the manner by which shared variance can be described as part of larger, related clusters. The clustering algorithm initially considers pairs of variables in terms of their similarity or “distance” (in arbitrary units). Linked pairs are then incorporated into larger clusters with the goal of minimizing a cost function (distance to be bridged), a process that can be thought of as minimizing the dissimilarity within clusters. As pairs become clusters, a cluster tree or dendrogram is created. The distance between neighbouring branches indicates the relative similarity of two measures, while advancing up the hierarchical cluster tree moves further away in terms of link distance, and therefore similarity.

Hierarchical models are useful as a descriptive tool for examining and visualising the structure of the dataset as a whole. However, they do not provide information as to the significance of any given cluster of behavioural measures. In contrast, exploratory factor analysis (EFA), which falls under the umbrella of structural equation modelling, can be used to formalise the relationships observed in the hierarchical models. This allows the researcher to establish the presence of underlying shared constructs via a number of fit statistics without applying a preconceived structure on the result.

For the EFA analysis, the smallest number of uncorrelated clusters that maximally explain the variance of the dataset was estimated. In this instance, parallel analysis with oblique rotation was employed to calculate this value. In a second step, the number of variables to be retained within the model was determined. A maximum likelihood estimation approach was applied, where variables that did not load significantly onto a particular factor or demonstrated significant cross loading (i.e. loaded onto more than one factor), were excluded from further testing. Finally, the model statistics were interrogated to formalize the shared variance across latent factors and the extent to which each variable contributed to its factor as a whole. The smallest number of factors that significantly explained the variance across the dataset was then accepted as the model of best fit. Model selection criteria included loading variables above 0.4 with no cross loading or freestanding variables, and significant X2/df ratio with Tucker-Lewis Index (TL-index) close to 1 and RMSEA < 0.06. Models were fit using Lavaan version 0.6-1 [E22] in R version 3.2.1 (R Core Team).

### Supplementary Results

**Supplementary Figure 1.**
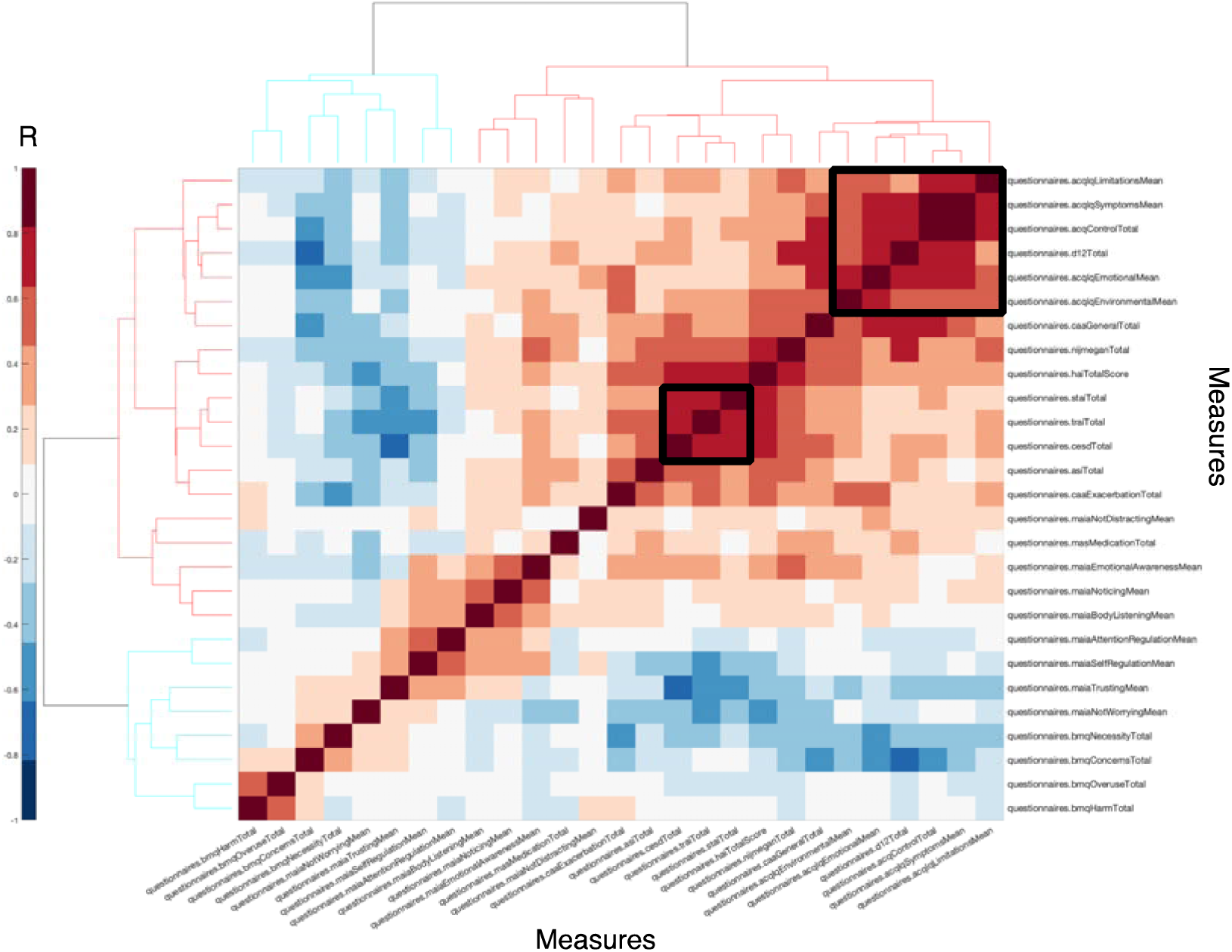
Clustergram of the questionnaire measures collected in the participants with asthma (n = 63). Questionnaire measures are ordered according to the strength of their correlation strength (R) so that any relationships between measures can be observed. Overlaid in black boxes are the two latent factors identified within the exploratory factor analysis.

**Supplementary Figure 2.**
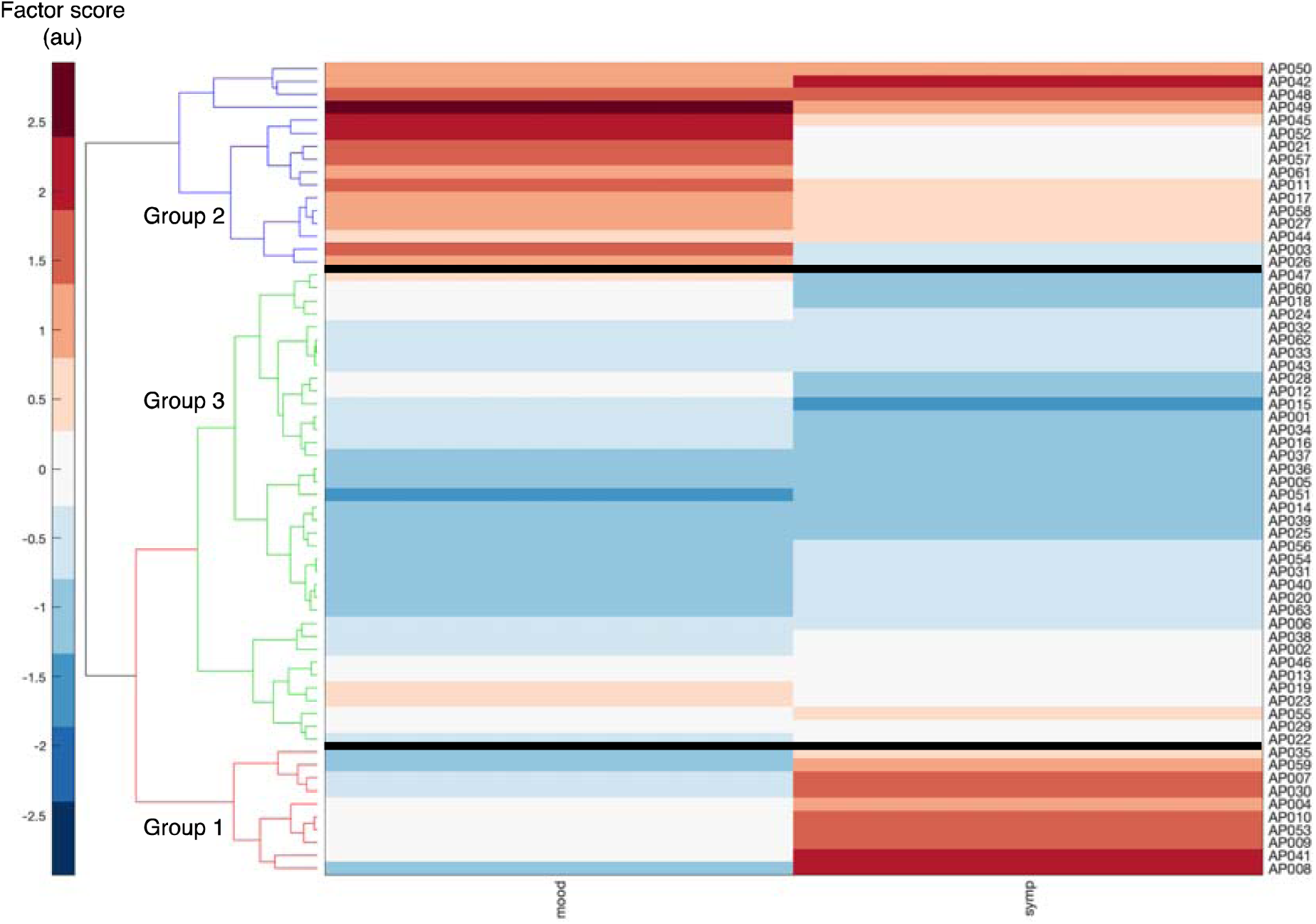
Clustergram of the asthma subject stratification, based on the first principle components of the latent questionnaire factors identified using exploratory factor analysis (‘Mood’ and ‘Symptoms’), where four group clusters can be observed.

**Supplementary Figure 3.**
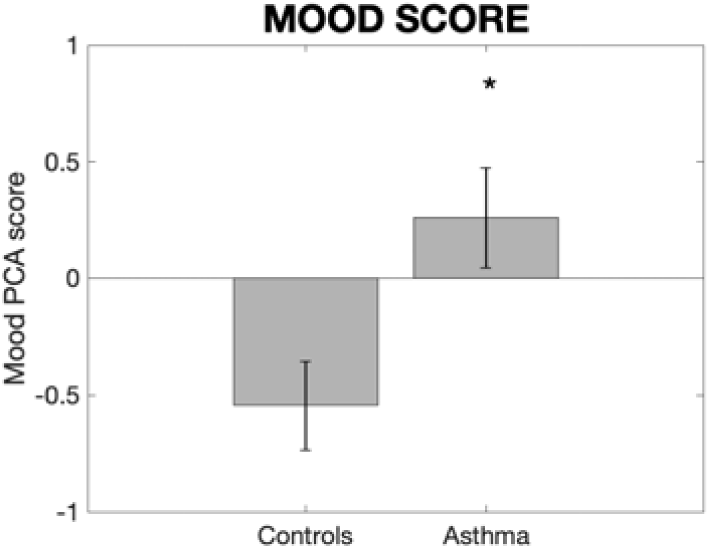
Group means and standard errors for the first principle component score of the ‘Mood’ factor (identified in the exploratory factor analysis) between individuals with asthma and healthy controls. A larger score denotes higher measures on the contributing questionnaires and thus worsened mood. * Significantly different from control group (p < 0.05, uncorrected).

**Supplementary Figure 4.**
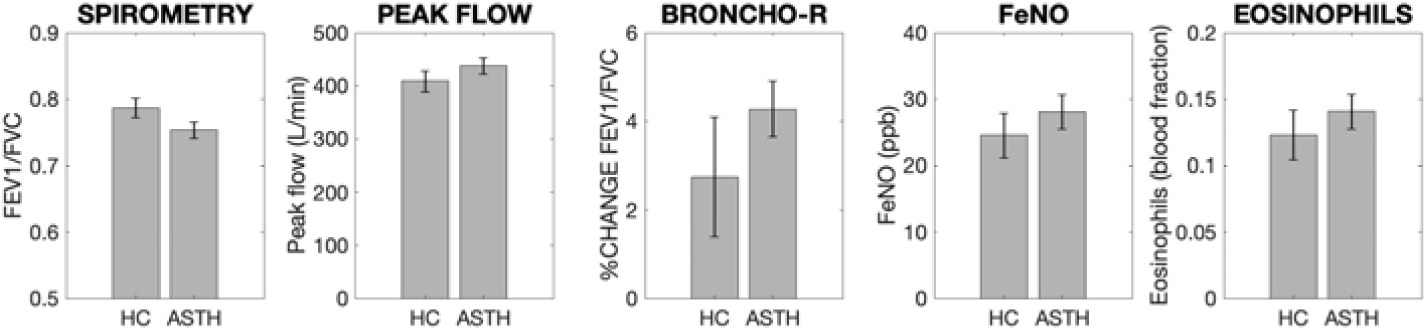
Group means and standard errors of the physiological measures for individuals with asthma (ASTH) and healthy controls (HC). No scores were significantly different between groups (significance was taken at p < 0.05, uncorrected). Abbreviations: BRONCHO-R, bronchodilator responsiveness; FeNO, fraction of exhaled nitric oxide.

**Supplementary Figure 5.**
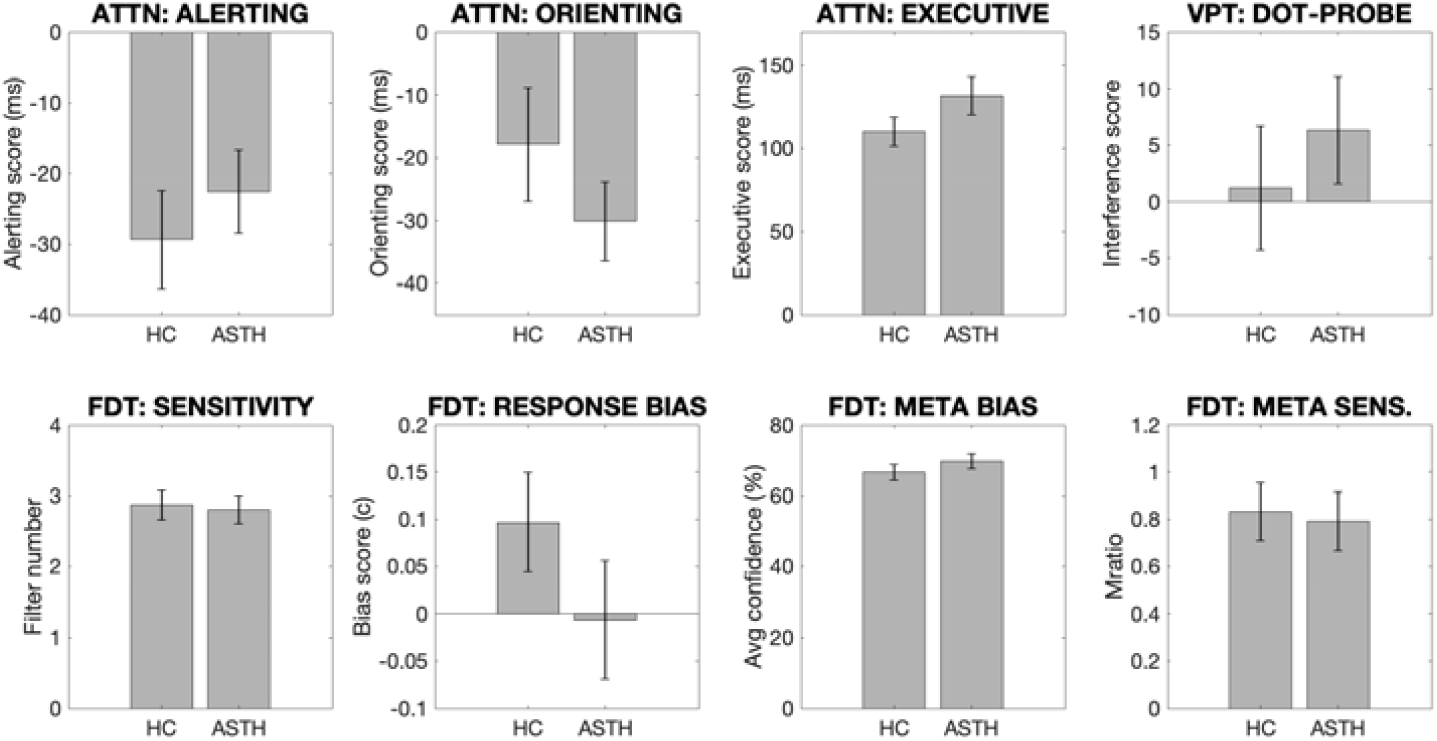
Group means and standard errors of the attention and interoceptive task measures for individuals with asthma (ASTH) and healthy controls (HC). No scores were significantly different between groups (significance was taken at p < 0.05, uncorrected). Abbreviations: ATTN, attention task; VPT, Visual Dot Probe Task; FDT, filter detection task; META BIAS, metacognitive bias; META SENS., metacognitive sensitivity.

**Supplementary Table 1.**
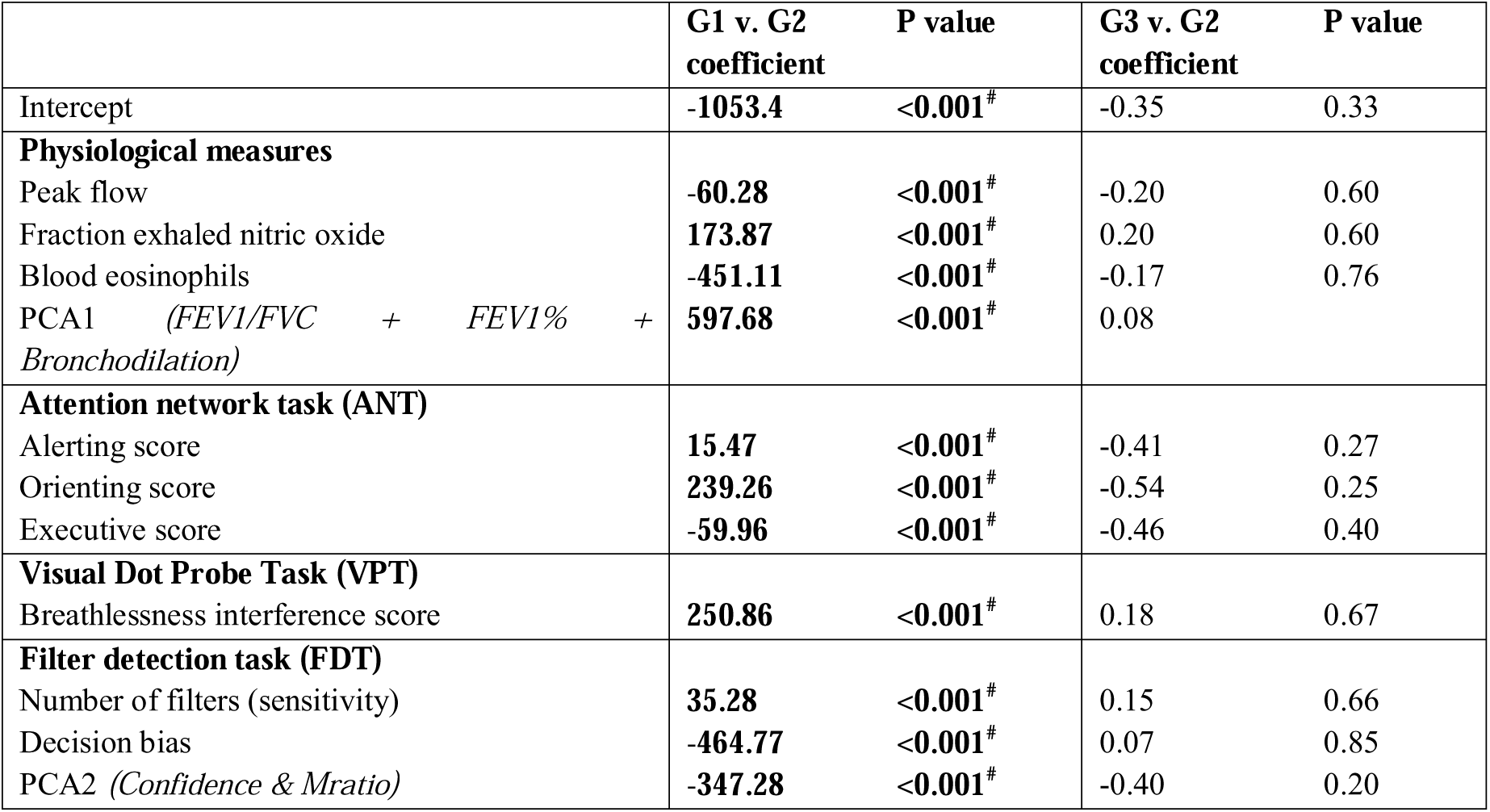
Logistic regression coefficients for asthma sub-group comparisons, where each of groups 1-3 were compared to group 4 (lowest mood and symptom scores). * Significant under the exploratory threshold of p < 0.05, and ^#^ Significant with FDR-correction for multiple comparison applied.

**Supplementary Table 2.** Missing data (reported as a percentage of data points) for each of the measures collected in both asthma and healthy controls.

|  | Asthma | Healthy controls |
| --- | --- | --- |
| Questionnaires | 0.3 | 1.1 |
| FEV1/FVC | 0 | 0 |
| FEV1 % predicted | 1.6 | 0 |
| Peak flow | 1.6 | 3.3 |
| Bronchodilator responsiveness | 0 | 0 |
| FeNO | 4.8 | 10 |
| Eosinophils | 11.1 | 10 |
| Attention task | 0 | 0 |
| Visual probe task | 0 | 0 |
| Filter detection task | 4.8 | 0 |

